# Expression of CD49f defines subsets of human regulatory T cells with divergent transcriptional landscape and function that correlate with ulcerative colitis disease activity

**DOI:** 10.1101/2021.02.22.432185

**Authors:** Harshi Weerakoon, Jasmin Straube, Katie Lineburg, Leanne Cooper, Steven Lane, Corey Smith, Saleh Alabbas, Jakob Begun, John J Miles, Michelle M Hill, Ailin Lepletier

**Affiliations:** Precision and Systems Biomedicine Laboratory, QIMR Berghofer Medical Research Institute, Herston, QLD, Australia; School of Biomedical Sciences, The University of Queensland, Brisbane, QLD, Australia; Department of Biochemistry, Faculty of Medicine and Allied Sciences, Rajarata University of Sri Lanka, Saliyapura, Sri Lanka; Gordon and Jessie Gilmour Leukaemia Research Laboratory, QIMR Berghofer Medical Research Institute, Herston, QLD, Australia; Translational and Human Immunology Laboratory, QIMR Berghofer Medical Research Institute, Herston, QLD, Australia; Inflammatory Bowel Diseases Research Group, Mater Research Institute, University of Queensland, Brisbane, QLD, Australia; School of Medicine, University of Queensland, Brisbane, Queensland, Australia; Mater Hospital Brisbane, Brisbane, QLD, Australia; Human Immunity Laboratory, QIMR Berghofer Medical Research Institute, Herston, QLD, Australia; Centre for Biodiscovery and Molecular Development of Therapeutics, James Cook University, Cairns, QLD, Australia; Centre for Clinical Research, Faculty of Medicine, The University of Queensland, Brisbane, QLD Australia; Laboratory of Vaccines for the Developing World, Institute for Glycomics, Southport, QLD, Australia

**Keywords:** adaptative immunity, cellular immunity, CD49f (integrin alpha 6), regulatory T cells, interleukin-17A (IL-17A), ulcerative colitis

## Abstract

**Objective:** Adoptive regulatory T cell (Treg) therapy is being trialled for treatment of different autoimmune disorders, including inflammatory bowel diseases (IBD). In-depth understanding of the biological variability of Treg in the human blood may be required to improve IBD immune monitoring and treatment strategies.

**Methods:** Through a combination of quantitative proteomic, multiparametric flow cytometry, RNA-sequencing data analysis and functional assays on Treg enriched from the blood of ulcerative colitis (UC) patients and healthy controls, we investigated the association between CD49f expression, Treg phenotype and function, and UC disease activity.

**Results:** High-dimensional analysis and filtering defined two distinct subsets of human Treg based on the presence or absence of CD49f with divergent transcriptional landscape and functional activities. CD49f negative Treg are enriched for functional Treg markers and present significantly increased suppressive capacity. In contrast, CD49f high Treg display a pro-inflammatory Th17-like phenotype and accumulate in the blood of UC patients. Dysregulation on CD49f Treg subsets in UC patients correlate with disease activity.

**Conclusion:** Overall our findings uncover the importance of CD49f expression on Treg in physiological immunity and in pathological autoimmunity.

## INTRODUCTION

Immune suppression through regulatory T cells (Treg) is pivotal for maintaining body homeostasis, controlling exaggerated immune responses against pathogens, and the prevention of immune cells attacking healthy tissue in the cases of autoimmunity, allergy, allograft rejection and foetal rejection during pregnancy (1). While the overall Treg cell population is defined as CD4^+^ T cells bearing a CD25^high^FoxP3^high^CD127^-^ phenotype, Treg found in peripheral circulation are highly heterogenic and have diverse function. At least 22 phenotypically different Treg subsets have been identified by mass cytometry in humans (2). Further, many activated conventional CD4^+^ T cells (conv CD4^+^) can also express CD25 and FoxP3 at low levels which cloud the specific identification of Treg (3). It is possible that a comprehensive multi ‘omic’ approach associating both proteomic and transcriptome analysis could lead more precise characterization of the various Treg subsets providing new insights into Treg mechanisms that guide homeostasis in health and dysfunction in disease.

Due to their multiple suppressive mechanisms, Treg represent a promising strategy for adoptive cell therapy for chronic inflammatory diseases. Treg are critical for commensal tolerance in the intestine, and a lack of intestinal tolerance can lead to chronic inflammation including inflammatory bowel diseases (IBD) consisting mainly of Crohn’s disease (CD) and ulcerative colitis (UC) (4–6). Evidence from both animal models and patients suggest that Treg therapy would be beneficial in the context of IBD. Treg have been expanded from patient’s blood and safely used in recent phase 1 studies designed for treatment of CD, type 1 diabetes mellitus, lupus and autoimmune hepatitis (7–9). Despite this great promise, the therapeutic use of Treg has been hampered by the biological variability of Treg populations in the peripheral blood. Effector Treg are heterogeneous and unstable following expansion; however, they do demonstrate increased suppressive function, higher efficacy and specificity in controlling immune responses compared with resting Treg (10).

Besides the loss of Treg suppressive function, infiltration of pro-inflammatory T-helper 17 (Th17) cells is also implicated in the pathogenesis of IBD (11). Interestingly, Treg differentiation is tightly linked to the development of Th17 cells, an effector T cell subset involved in the induction of inflammation and implicated in autoimmune tissue injury through the production of interleukin 17A (IL-17A) and other pro-inflammatory cytokines (12). Whereas both the induction of peripheral Treg from resting CD4^+^ T cells and the maintenance and function of natural Treg are dependent on transforming growth factor beta (TGF-β) signaling, the presence of IL-6 inhibits TGF-β-mediated FoxP3 induction and drives cells towards a Th17 phenotype. A subset of Treg cells expressing the Th17-associated markers retinoid-related orphan receptor-gamma t (RORγt) and chemokine receptor 6 (CCR6), in addition to FoxP3, have also been reported *in vivo* and is increased in the intestinal mucosa and among peripheral blood mononuclear cells (PBMC) circulating in IBD patients compared with healthy controls (12–14). However, the mechanisms that underpin the development of these Th17-like Treg cells are still under debate due to the high Treg cell plasticity, which can be detrimental in the setting of autoimmune diseases.

Several T cell subsets express adhesion receptors known as integrins, such as CD49a, CD49b, CD49d and CD49f that have been reported to modulate various aspects of T cell biology including cell differentiation, migration and functionality (15–18). It is possible that CD49f (integrin alpha 6) expression on CD4^+^ T cells is associated with the pathogenesis of IBD, as CD49f is increased on the surface of circulating conv CD4^+^ cells that migrate out of the colonic mucosa of patients with active IBD (19).

In order to assess the impact of CD49f expression on Treg-mediated immune responses in health and disease, we investigated the association between CD49f expression, Treg phenotype and function, and clinical outcomes in IBD patients. Comparative proteomics between Treg and conv CD4^+^ cells reveal CD49f to be divergently expressed among circulating Treg. Using high-dimensional analysis and filtering, we define two subsets of Treg, based on the presence or absence of CD49f, with divergent transcriptional landscape and functional activities. Our data reveal that CD49f negative (CD49f neg) Treg exert high suppression on conv CD4^+^ cell proliferation, associated with elevated expression of FoxP3 and the immune checkpoint receptors, CD39 and CTLA4. In contrast, CD49f high Treg produce abnormal levels of IL-17A under TCR-mediated activation, concurrently expressing higher levels of CCR6, and are increased in PBMC of UC patients compared with healthy controls. Notably, an elevated CD49f high/CD49f negative effector Treg ratio (CD49f^eR^) in patients’ blood is a predictor of active disease in UC. Taken together, our findings demonstrate that CD49f expression on Treg impacts human physiological immunity and influence the development of IBD and possibly other autoimmune disorders.

## RESULTS

### CD49f is divergently expressed among human regulatory T cells

Treg cells are generally identified as a CD4^+^ T cell subset with suppressive properties and a high phenotypic and functional diversity (2). To allow better Treg characterization in humans, we set out to identify Treg-enriched cell surface proteins using comparative proteomics between regulatory T cells (Treg, CD4^+^CD25^high^CD127^-^) and conv CD4^+^ cells (CD4^+^CD25^-^). Treg with high purity were obtained from human PBMC through sequential magnetic and flow cytometry cell sorting (Figure 1a). As expected, purified Treg showed high expression of FoxP3, proportional to their CD25 levels, whereas FoxP3 expression on conv CD4^+^ cells was similar to the unstained controls (Figure 1a). To identify proteins preferentially enriched in Treg compared to conv CD4^+^ cells, we conducted label-free quantitative proteomics using data dependent acquisition (DDA-MS). Inspection of the maxLFQ normalized intensity values showed missing value in <10% of proteins in each sample (Supplementary figure 1a), confirming the acquisition of high quality DDA-MS data for unambiguous label free quantification. We identified a total of 4,177 protein groups at 1% of false discovery rate (FDR) (Supplementary figure 1b). 2,788 proteins were quantified using single UniProt accessions with at least 2 unique and razor peptides in more than 50% of the samples and thus selected for differential expression (DE) analysis (Supplementary figure 1b). Most of these proteins were quantified based on intensities of more than 5 peptides (Supplementary figure 1c) and showed distribution pattern common to all the samples analyzed (Supplementary figure 1d). Hierarchical cluster analysis based on Euclidian distance clearly separated the proteomic data into two groups according to the CD4^+^ T cell subset analyzed (Supplementary figure 1e). This was further confirmed by principal component analysis (PCA), in which two clear clusters were observed in the first principal component (Figure 1b). Most of the proteins within each subset had < 2% co-variability (Supplementary figure 1f), verifying the consistency and reproducibility of the obtained label free quantitative DDA-MS data. In addition, we used Ingenuity Pathway Analysis (IPA, Qiagen bioinformatics, USA) to identify the subcellular distribution of these identified proteins. As expected, most of the detected proteins from whole cell lysates were derived from the cytoplasm and nucleus whereas 180 proteins (∼ 6% of the total quantified proteins) were annotated as plasma membrane proteins and considered of interest as potential uncharacterized Treg surface markers (Figure 1c and Supplementary table 1). Statistical analysis identified 227 proteins as DE (FDR < 0.05 and log2FC >1 or <-1) between donor matched Treg and conv CD4^+^ cells and indicated that only 10% of the global Treg proteome was significantly different from conv CD4^+^ (Supplementary table 2). Of the DE proteins, 72% (n = 166) were up regulated in Treg cells, including FoxP3 with log2 fold change of 6.29 (Figure 1d). As a candidate Treg surface marker, we selected the plasma membrane protein CD49f, which showed a 3.12 log2 fold increase in relation to conv CD4^+^ (Figure 1d). Subsequent flow cytometric validation using a CD49f monoclonal antibody revealed that CD49f is heterogeneously expressed in human Treg, allowing the identification of 3 distinct Treg populations characterized as CD49f neg, dim and high cells (Figure 1e). In accordance with the proteomic data, both CD49f mean fluorescence intensity (MFI) and the fraction of CD49f high cells were significantly increased in human Treg when compared to conv CD4^+^ cells (Figure 1f).

**Figure 1.**
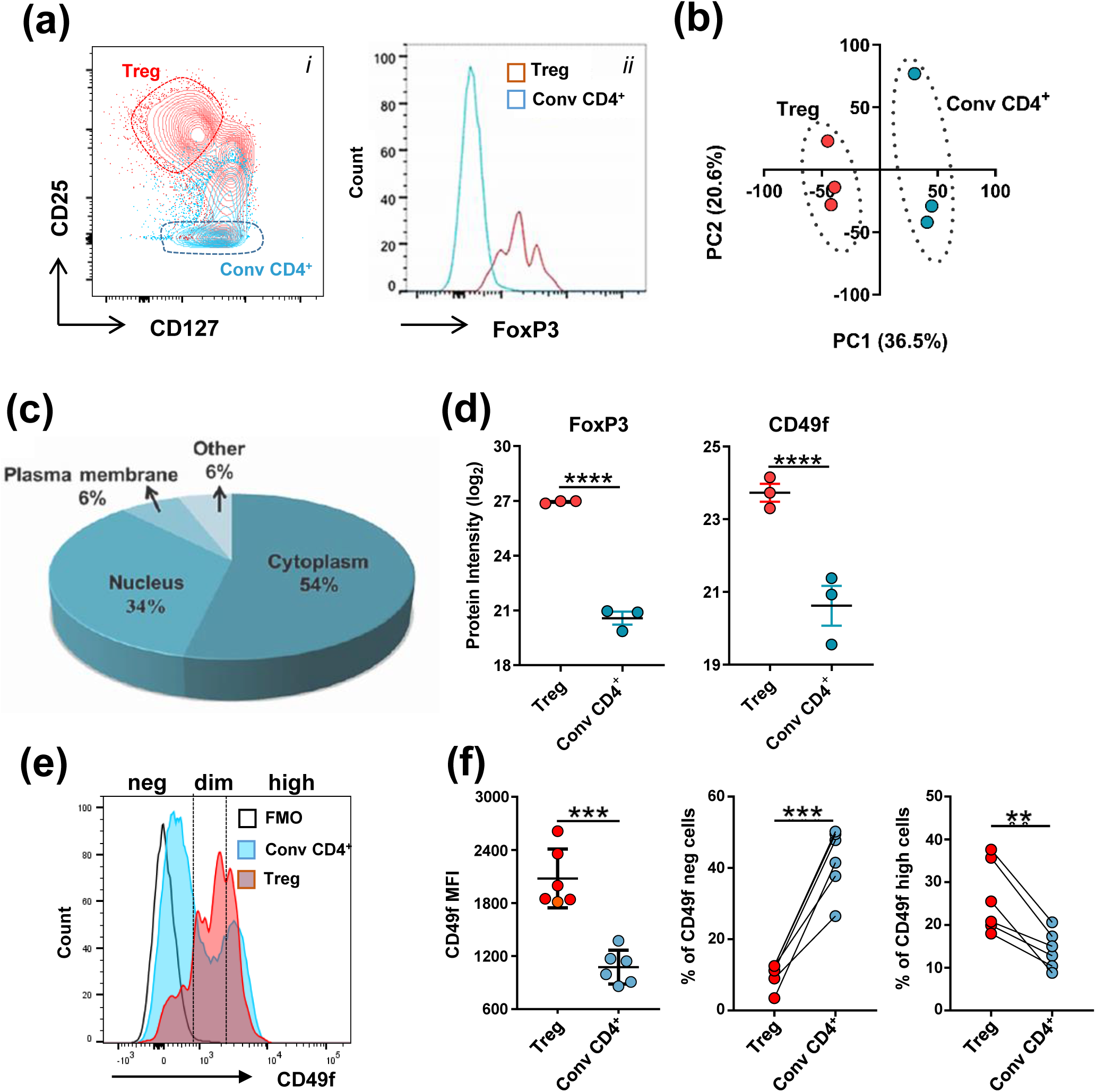
CD49f is divergently expressed among human regulatory T cells. Treg (CD4^+^CD25^hi^CD127^-^) and conv CD4^+^ T cells (CD4^+^CD25^-^) were isolated from fresh PBMC of healthy individuals for proteomic and flow cytometric characterization. (a) Gating strategy for cell sorting of Treg and conv CD4^+^ cells. *i* Contour plots in red show positively enriched CD4^+^CD25^+^ T cells by magnetic sorting and conv CD4^+^ cells in the negative fraction are shown in blue. The dotted line represents the populations subsequently enriched by FACS sorting. *ii* FoxP3 staining of enriched Treg and conv CD4^+^ cells for confirmation of sort purity. (b) Principal component (PC) projections of individual Treg and conv CD4^+^ cells obtained from proteomic analysis of three healthy controls. PC1 (36.5% variance) and PC2 (20.6% variance) are shown. (c) Pie chart displaying the subcellular distribution of quantified proteins using IPA. (d) Proteomic profiles for FoxP3 and CD49f expression on enriched Treg and conv CD4^+^ cells. **** FDR < 0.0001. Multiple t-test. Bars represent standard error of mean. (e) Overlaid histograms representing CD49f MFI on Treg and conv CD4^+^ cells by flow cytometric analysis. Cell subsets were defined based on CD49f intensity as negative (neg), dim or high cells. A fluorescence minus one (FMO) control was used to normalize protein expression. (f) CD49f MFI and fraction of CD49f neg and CD49f high cells in Treg and conv CD4^+^ subsets. Data was obtained from six healthy controls. ***P<0.01, ***P<0.001. Non-parametric paired t-test.

Thus, CD49f might define a unique subset of Treg with unexplored functions.

### CD49f impacts Treg immunosuppressive ability and IL-17A production

We next wanted to understand the effect of CD49f expression on Treg function. CD49f high and CD49f neg Treg were sorted from PBMC of healthy volunteers (Figure 2a) and their activity was measured by *in vitro* suppression assay of autologous conv CD4^+^ cell proliferation in the presence of OKT3 antibodies (1μg/ml) and irradiated allogenic PBMC. Stimulated conv CD4^+^ cells without Treg were cultured in the same assay for definition of appropriate controls (Figure 2b). An increased proliferation of conv CD4^+^ cells was observed when the cells were co-cultured in the presence of CD49f high Treg when compared to CD49f neg controls. This effect was detected across the multiple Treg: conv CD4^+^ cell ratios analyzed (Figure 2b and 2c). CD49f high Treg showed a suppressive potential similar to total Treg, which was detected only when cells were cultured in a Treg: conv CD4^+^ cell ratio below 1:8 (Figure 2c, Supplementary figure 2a). Contrarily, CD49f neg Treg were highly suppressive even when cells were cultured in a Treg: conv CD4^+^ cell ratio of 1:16 (Figure 2c). CD49f neg Treg purified from five different donors consistently presented increased ability to suppress CD4^+^ T cells proliferation, averaging 65.8 ± 6.89% versus 49.4 ± 3.37% of suppression observed in the CD49f high fraction (Figure 2d).

**Figure 2.**
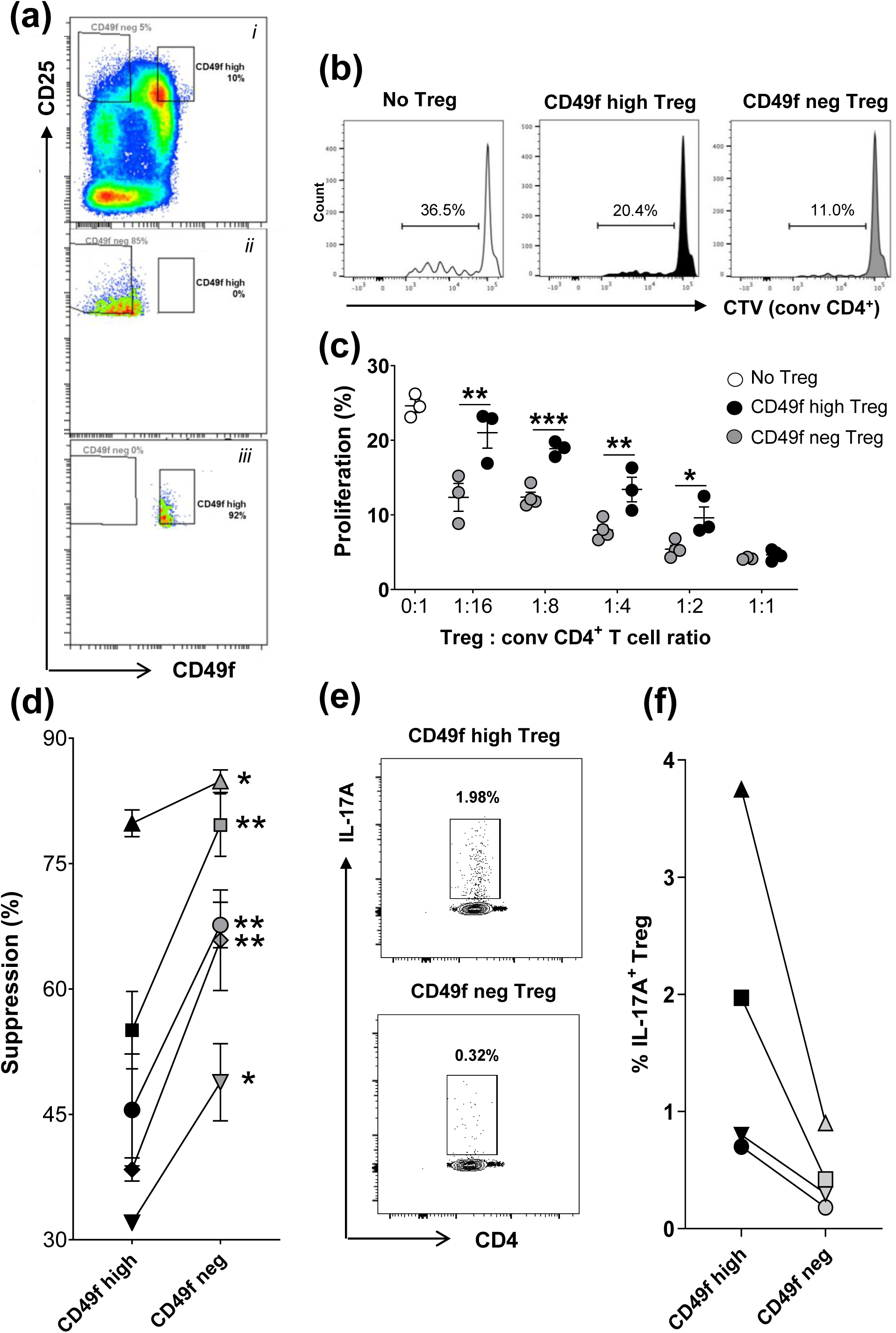
CD49f impacts Treg immunosuppressive ability and IL-17A production. (a) *i,* Gating strategy used to purify subsets of CD49f high and CD49f neg Treg (CD4^+^CD127^-^CD25^hi^ cells) from human PBMC by FACS sorting. Post-sorting check of *ii,* CD49f neg and *iii,* CD49f high in enriched subsets. (b) Histograms representing the fraction of proliferating (CTV^-/low^) conv CD4^+^ cells cultured in the presence of CD49f neg, CD49f high Treg (1 Treg:8 conv CD4^+^ cells) or in the absence of Treg. Conv CD4^+^ cells were loaded with CTV and activated with CD3/CD28 beads in the presence of CD49f high or CD49f neg Treg cells during five days of *in vitro* culture. (c) Graph showing the fraction of proliferative conv CD4^+^ cells cultured in the presence or absence of Treg CD49f high or CD49f neg cells at various Treg: conv CD4^+^ cell ratios. Data represents one donor analyzed and includes 3-4 technical replicates. *P<0.05, **P<0.01 and ***P<0.001 CD49f high versus CD49f neg Treg. Non-parametric paired t-test. Bars represent standard error of mean. (d) Fraction of CD49f high and CD49f neg Treg mediated suppression on conv CD4^+^ cell proliferation calculated as 100−((proliferated cells with Treg/proliferated cells with no Treg) x 100). Data was obtained from five donors analyzed and includes 3-4 technical replicates. *P<0.05 and **P<0.01 CD49f high versus CD49f neg Treg. Non-parametric paired t-test. Bars represent standard error of mean. (e) Fraction of IL-17A^+^ cells in CD49f neg and CD49f high Treg following overnight activation in the presence of CD3/CD28 Dynabeads and recombinant IL-2 during five days of *in vitro* culture. Data was obtained from four healthy controls analyzed. (f) FACS plot representing IL-17A production by CD49f high and neg Treg cell fractions.

Based on previous studies indicating the existence of Treg that have the capacity to produce pro-inflammatory cytokines whilst retaining FoxP3 expression (12, 14), we sought to investigate an association between interleukin-17A (IL-17A) and interferon gamma (IFNγ) production by Treg and CD49f expression. Enriched CD49f high and CD49f neg Treg were activated with CD3/CD28 Dynabeads in the presence of recombinant human IL-2 and brefeldin A and analyzed by flow cytometry for intracellular expression of IL-17A and IFNγ. In all donors evaluated, the proportion of IL-17A^+^ cells was three to five-fold higher in CD49f high cells compared to their CD49f neg control population, comprising 1.8 ± 0.7% and 0.45 ± 0.16% of the cells analyzed cells, respectively (Figure 2e and f). Similar to CD49f neg Treg, 0.6% of total Treg expressed IL-17A in the same experiment (Supplementary figure 2b). No association between CD49f expression and IFNγ was observed in activated Treg (Supplementary figure 2c) and most of IL-17A^+^ cells did not co-expressed IFNγ (Supplementary figure 2d). Thus, IL-17A-expressing Treg might represent a distinct population of cytokines-producing cells defined by CD49f expression.

Taken together, our data show that CD49f expression impacts Treg immunosuppressive abilities and IL-17A production.

### RNA-sequencing uncovers distinct subsets of regulatory T cells defined by CD49f expression

To comprehensively profile the relevant immune pathways associated with CD49f high and CD49f neg Treg, we performed next generation RNA sequencing (RNA-Seq) on the two populations using high purity flow cytometry sorting of Treg from the peripheral blood of healthy individuals. We compared the transcriptional profiles between the sorted subsets, which revealed two distinct Treg populations by PCA, in which two clear clusters were observed in the first principal component (Figure 3a). Transcriptional differences between CD49f neg and CD49f high cells were further confirmed by hierarchical cluster analysis (Figure 3b). From the 17,936 genes identified, 668 genes were DE in donor matched CD49f high and CD49f neg cells (FDR < 0.05, absolute log2FC>1) (Supplementary figure 3a, Supplementary table 3). In CD49f high Treg, 225 genes were upregulated in comparison to their negative controls (Supplementary figure 3b). Gene expression differences were found in key molecules relevant for Treg effector function. CD49f neg Treg presented increased expression of classical Treg markers including *CTLA4, ENTPD1 (CD39), ICOS, LAG3* and *FOXP3* when compared to CD49f high controls. In contrast, CD49f high Treg expressed higher levels of genes associated with Th17 effector cytokine signaling, including *RORC, CCR6, RORA, GPR65* and *MYC* (Figure 3c). Consistently, gene set enrichment analysis (GSEA) of a gene list derived from comparing Treg and CD4^+^ Th17 cells in healthy human PBMC (GSE107011), revealed that CD49f high Treg gene expression profile corresponded to the Th17 signature while genes upregulated in CD49f neg Treg overlapped with classical Treg gene signature (Figure 3d).

**Figure 3.**
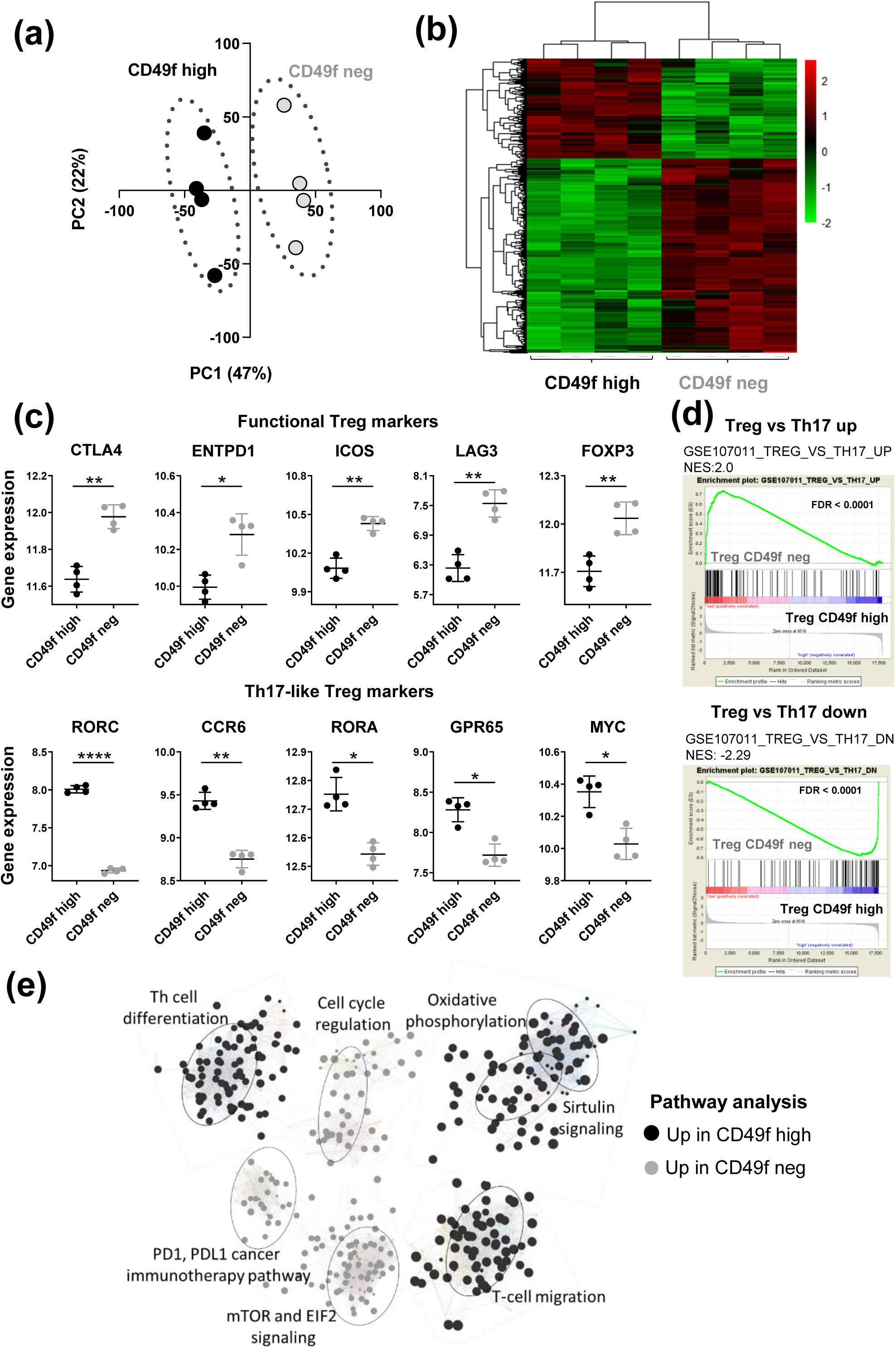
RNA-sequencing uncovers distinct subsets of regulatory T cells defined by CD49f expression. Gene list obtained from RNA-Seq data analysis of CD49f high and CD49f neg Treg sorted from PBMC of four healthy controls. (a) Principal component (PC) projections of individual Treg and conv CD4^+^ cells obtained from RNA-Seq analysis of four healthy controls. PC1 (47% variance) and PC2 (22% variance) are shown. (b) Heatmap analysis of differentially expressed genes from comparing CD49f high and CD49f neg Treg. Each column represents individual donors across genes differentially expressed between CD49f high versus CD49f neg Treg (n=4; FDR < 0.01). (c) Graphs showing DE genes in CD49f neg *versus* CD49f high Treg. Genes were grouped as functional Treg markers, associated with immunosuppression, or IL-17-like Treg markers, associated with Th17 signature. P*<0.05, P**<0.01, P***<0.0005. Non-parametric t test. Bars represent standard error of mean. (d) Gene set enrichment analysis of Treg and Th17 cells in healthy human PBMC (GSE107011). Data demonstrate overlap between genes expressed on CD49f neg Treg and genes upregulated on classical Treg (upper graph; FDR < 0.001), and between genes expressed on CD49f high Treg and genes upregulated in Th17 cells (bottom graph, FDR < 0.001). (e) Top seven pathways characterizing the differences between CD49f neg and CD49f high Treg gene expression programs identified by IPA. Nodes denote genes composing the pathways in the IPA database and upregulated in CD49f neg (grey) or CD49f high T regs (black). Lines show the connectivity between nodes and the node size indicates their degree of connectivity.

To identify altered pathways activity in these Treg subsets, DE genes were analyzed using IPA core analysis. Pathways upregulated in CD49f high Treg were those involved in the oxidative metabolic pathway, cell migration, sirtulin signaling and T helper cell differentiation. In comparison, CD49f neg Treg were enriched for genes associated with cell cycle regulation, immune checkpoint modulation and mTOR and EIF2 signaling, which are critical regulators of Treg homeostasis and function (Figure 3e).

These data further validate the functional divergence between CD49f high and CD49f neg Treg biology via significant transcriptional differences.

### CD49f is associated with divergent effector regulatory phenotype and function in Treg

Next, we investigated the impact of CD49f expression on the different subsets of Treg cells. As previously described, the simultaneous assessment of FoxP3 and CD45RA allows for the identification of three distinct subpopulations of human FoxP3-expressing CD4^+^ T cells: resting Treg (CD45RA^+^FoxP3^low^), effector Treg (CD45RA^−^FoxP3^high^), and FoxP3^+^ non Treg cells (CD45RA^−^FoxP3^low^), which produce pro-inflammatory cytokines and lack suppressive capacity (20, 21). In this aim, we sought to validate DE genes originally identified by RNA-Seq using flow cytometric analysis of effector Treg, resting Treg and FoxP3^+^ non Treg cells (Figure 4a). As expected, CD4 MFI was similar among the three subsets analyzed but the MFI for CD25, FoxP3, CD39, CTLA4 and CCR6 was increased in effector cells in relation to both resting Treg and FoxP3^+^ non Treg (Supplementary figure 4). Whereas CD49f fluorescence intensity did not differ between effector and resting cells (Figure 4b), CD49f specifically impacted the phenotype of effector Treg. In accordance with the RNA-Seq data, we observed an increased MFI for CD39, CTLA4 and FoxP3 expression in CD49f neg versus CD49f high effector cells (Figure 4c and 4d). In contrast, CCR6 MFI directly correlated with the level of CD49f expression on effector Treg (Figure 4c and 4d). CD49f expression was also associated with MFI for CTLA4, CD39 and CCR6 in FoxP3^+^ non Treg (Figure 4c). CD49f dim cells expressed intermediate levels of each marker quantitated in both effector and FoxP3^+^ non Treg cells. Resting Treg expressed similar levels of CD39, CTLA4, FoxP3 and CCR6 to FoxP3^+^ non Treg, which did not correlate with CD49f MFI (Supplementary figure 4 and Figure 4c).

**Figure 4.**
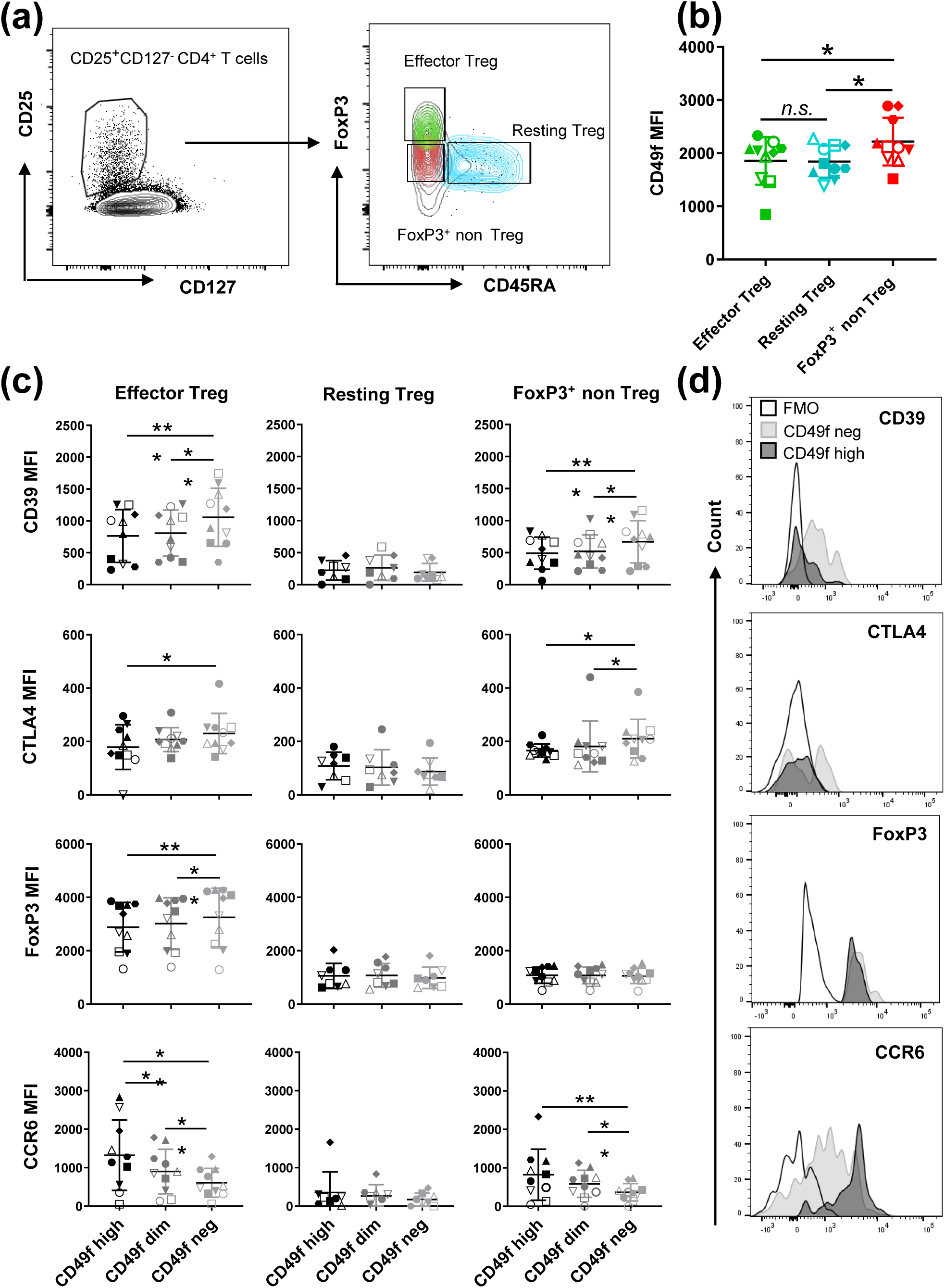
CD49f correlates with divergent effector regulatory phenotype and function in Treg. Flow cytometric analysis of Treg subsets in PBMC from ten healthy controls. (a) Representative gating strategy of Treg cells (CD4^+^CD25^+^CD127^-^), classified as effector Treg (FoxP3^high^CD45RA^-^), resting Treg (FoxP3^low^CD45RA^+^) and non-Treg cells FoxP3^+^ (FoxP3^low^CD45RA^-^). (b) CD49f MFI across Treg subsets defined by FoxP3 and CD45RA expression. P*<0.05, *n.s.* = non-significant. Non-parametric one-way ANOVA test with Bonferroni correction. Bars represent standard error of mean. (c) Graphs representing the correlation between CD49f expression and the MFI of the functional Treg markers CD39, CTLA4, FoxP3 and the Th17-associated chemokine receptor CCR6 in Treg subsets. Each symbol represents an individual donor analyzed. P*<0.05, P**<0.01, P***<0.0005. Non-parametric one-way ANOVA test with Bonferroni correction. Bars represent standard error of mean. (d) Histograms represent expression profile of functional markers and CCR6 across CD49f high and CD49f neg effector Treg. A fluorescence minus one (FMO) control was used to normalize protein expression.

Thus, combined validation using different platforms suggests that CD49f impacts Treg function and is a lead target for Treg investigation.

### CD49f expression on effector regulatory T cells correlates with disease activity in UC patients

Because CD49f expression on effector Treg could potentially impact autoimmune diseases in which Treg play a unique role, and CD49f has been reported to modulate CD4^+^ T cell homing during IBD (19), we hypothesized that CD49f expression on circulating human Treg may be altered in autoimmune conditions, such as UC. To evaluate this, we characterized CD49f expression using flow cytometry in circulating Treg from a cohort of UC patients who presented active or non-active disease at time of sampling (Table 1) and age matched volunteer healthy controls. We noticed a trend towards reduction of total Treg in UC patients (Figure 5a). While conv CD4^+^ cells were reduced in UC patients with active disease when compared to healthy controls, the fraction of Treg did not associate with UC disease activity (Supplementary figure 5a and 5b). Interestingly, CD49f was significantly enriched in the effector Treg subset from UC patients’ blood when compared with healthy controls (Figure 5b and 5c). A corresponding decrease in CD49f neg effector Treg was detected in UC patients with active disease in relation to healthy controls (Figure 5d). Notably, the ratio of CD49f high/CD49f negative effector Treg (CD49f^eR^) in the peripheral blood significantly correlated with UC disease activity (R = 0.275; P = 0.004) (Figure 5e and 5f). A minor association between CD49f expression and UC disease activity was observed in resting Treg (Supplementary figure 5c-5e).

**Figure 5.**
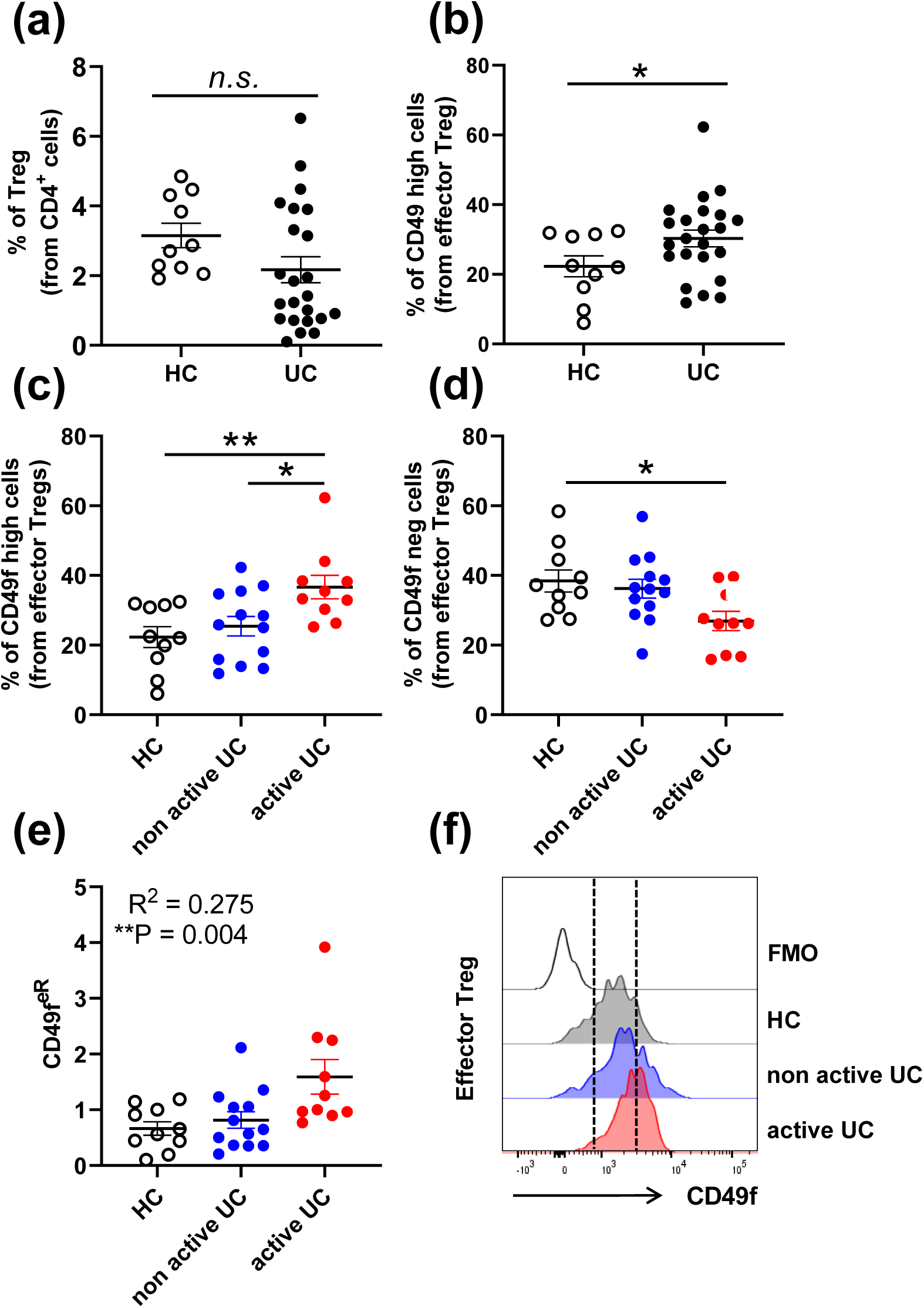
CD49f expression on effector regulatory T cells correlates with disease activity in UC patients. (a) Frequencies of Treg in PBMC from patients with ulcerative colitis (UC, n =23) and healthy control (HC, n = 10). N.s. = non-significant. Non-parametric t-test. Bars represent standard error of mean. (b) Proportion of CD49f high effector Treg in PBMC isolated from UC patients and healthy controls. P*<0.05. Non-parametric t-test. Bars represent standard error of mean. (c) Proportion of CD49f high cells in effector Treg. P*<0.05, P**<0.01. Non-parametric one-way ANOVA test with Bonferroni correction. Bars represent standard error of mean. (d) Proportion of CD49f neg cells in effector Treg. P*<0.05, P**<0.01. Non-parametric one-way ANOVA test with Bonferroni correction. Bars represent standard error of mean. (e) Data summary of the ratio of CD49f high effector Treg to CD49f neg effector Treg (CD49f^eR^, R^2^=0.275; P = 0.004). (f) Representative histograms showing CD49f distribution across effector Treg cells characterized in UC patients with active and non-active disease, and healthy controls (HC). A fluorescence minus one (FMO) control was used to normalize protein expression.

**Table 1.**
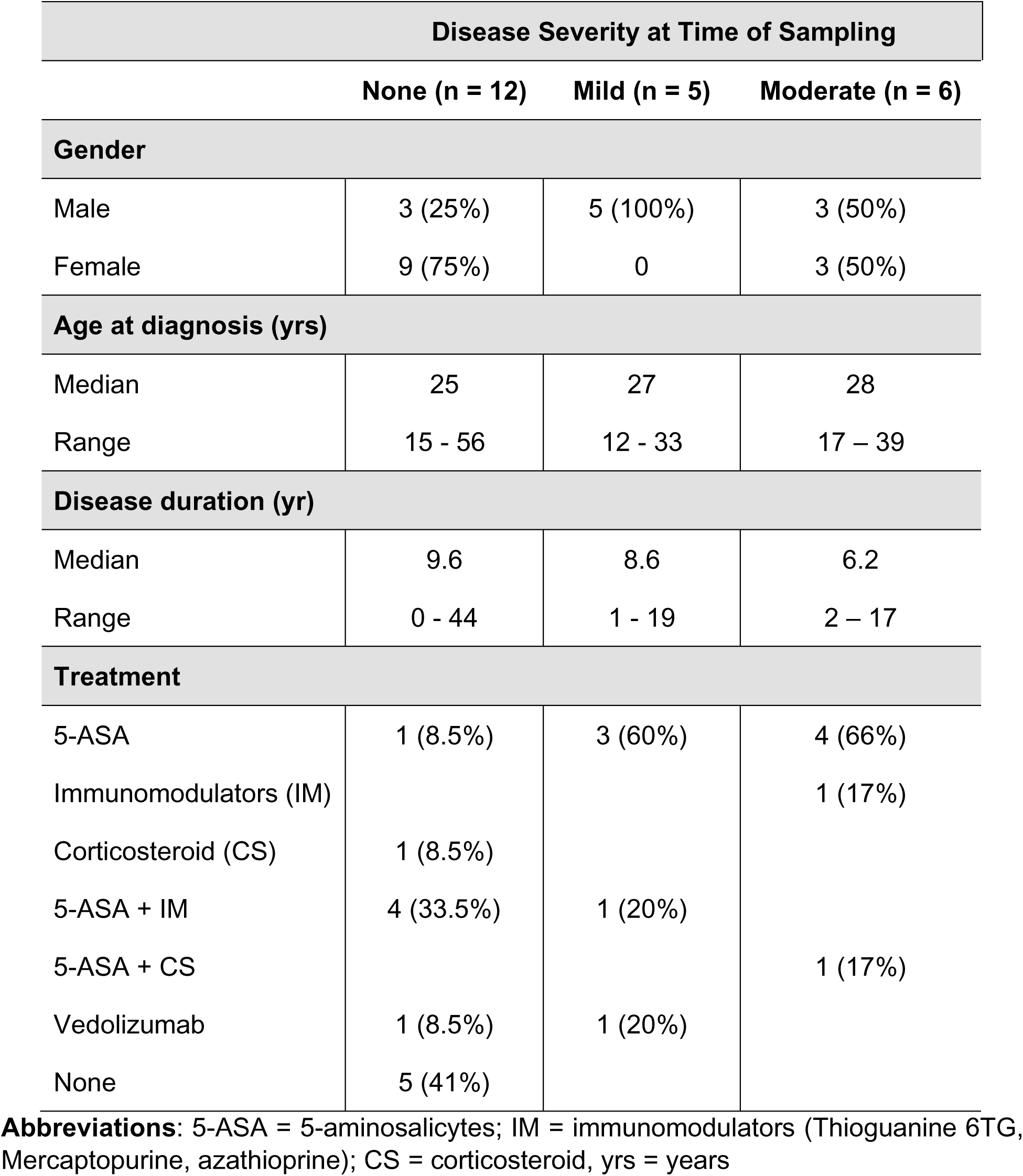
Demographics and clinical data of the ulcerative colitis study population.

| Disease Severity at Time of Sampling |  |  |  |
| --- | --- | --- | --- |
|  | None (n = 12) | Mild (n = 5) | Moderate (n = 6) |
| **Gender** |  |  |  |
| Male | 3 (25%) | 5 (100%) | 3 (50%) |
| Female | 9 (75%) | 0 | 3 (50%) |
| **Age at diagnosis (yrs)** |  |  |  |
| Median | 25 | 27 | 28 |
| Range | 15 - 56 | 12 - 33 | 17 – 39 |
| **Disease duration (yr)** |  |  |  |
| Median | 9.6 | 8.6 | 6.2 |
| Range | 0 - 44 | 1 - 19 | 2 – 17 |
| **Treatment** |  |  |  |
| 5-ASA | 1 (8.5%) | 3 (60%) | 4 (66%) |
| Immunomodulators (IM) |  |  | 1 (17%) |
| Corticosteroid (CS) | 1 (8.5%) |  |  |
| 5-ASA + IM | 4 (33.5%) | 1 (20%) |  |
| 5-ASA + CS |  |  | 1 (17%) |
| Vedolizumab | 1 (8.5%) | 1 (20%) |  |
| None | 5 (41%) |  |  |

**Abbreviations:** 5-ASA = 5-aminosalicylates; IM = immunomodulators (Thioguanine 6TG, Mercaptopurine, azathioprine); CS = corticosteroid, yrs = years

In summary, our data support the notion that active UC is associated with increased CD49f expression on circulating Treg and that assessment of CD49f ratios within the effector Treg compartment may be a useful predictor of disease activity.

## DISCUSSION

Through a combination of *in vitro* functional studies, quantitative proteomics, transcriptome deep sequencing and phenotypic analyses, we identified unexplored subsets of Treg defined by CD49f expression. This finding addresses a gap in understanding Treg immunomodulatory function in homeostasis and human diseases. The CD49f neg Treg subset exhibits a unique phenotypic profile with significantly increased suppressive capacity. In contrast, the frequencies of Th17-like CD49f high Treg correlated with the activity of UC, suggesting that subset exclusion based on CD49f high expression on Treg may constitute a promising strategy to maximize the efficacy and safety of Treg-based immunotherapy for treating IBD patients. Understanding the pathogenic role of CD49f high Treg in inflammatory disorders may provide insight into the drivers of maladaptive inflammation.

CD49f plays an important and conserved role in stem cell biology (22). It belongs to the integrin family of receptors, which are structurally characterized as transmembrane adhesion receptors that mediate cell–cell and cell–extracellular matrix adhesion and induces bidirectional signaling across the cell membrane that regulate proliferation, activation, migration and homeostasis (22). CD49f often dimerizes with β1 and β6 integrins to form heterodimers including α6β1 and α6β4 that act as primary receptors for laminins present in their niche (23). Accumulation of evidence from human and mouse models show that defects in integrin expression or unintentional inflammation against healthy host tissue result in serious immunodeficiency and many autoimmune conditions (24, 25). Accordingly, integrins such as CD49b and CD49d have been recently described to modulate various aspects of Treg biology (17, 18, 26, 27). Human CD49d^-^ Treg have been reported to present enhanced immunosuppressive function than their CD49d^+^ counterparts (18). This finding is supported by more recent evidences that CD49d^dim/-^ expression enriches for a subset of cells with suppressive capacity within the CD8^+^CD122^+^ population of effector T cells (28). In contrast, the expression of CD49b on mature Treg that survey the skin and vascular tissues resulted in superior suppressive capacity and decreased disease severity in a mouse model of T cell-induced arthritis, partially dependent on IL-10 secretion (27, 29). Unlike CD49d and CD49b, information on the Treg immune modulating capacity by CD49f is scarce.

In this study, we show that negative expression of CD49f defines a subset of effector Treg in the human blood with high suppressive capacity and increased expression of the functional Treg markers CTLA4, CD39 and FoxP3. FoxP3 expression is not only associated with Treg differentiation towards a suppressive phenotype but also a prerequisite for stabilizing the Treg lineage (30). Similarly, high expression of CD39 on human Treg drives cell stability and function under inflammatory conditions through the conversion of ATP into adenosine and AMP (31), whereas deficiency of CTLA4 in Treg is associated with development of spontaneous systemic lymphoproliferation and fatal T cell-mediated autoimmune disease (32). Interestingly, although CD49f expression on resting and effector Treg appear similar, CD49f only impacted the expression of functional markers on effector Treg, suggesting that rather being a marker for effector compared with resting Treg, CD49f is associated with suppressive function and IL-17A expression in the effector compartment.

UC is a chronic autoimmune disease characterized by infiltration of inflammatory cells into the lamina propria of the intestinal tract. Various subsets of intestinal lamina propria T cells are believed to traffic, via the systemic circulation, from gut-associated lymphoid tissue (33). Even though it is assumed that absence of Treg leads to IBD in both human and mice models (4–6), there is little evidence to suggest that IBD patients simply lack Treg in the circulation and/or the affected tissues (34–36). In our cohort, UC patients had a lower number of circulating Treg, but this was not statistically significant. However, CD49f high effector Treg were significantly increased in UC patients in comparison to healthy controls, and an increased ratio of CD49f high/CD49f negative effector Treg (CD49f^eR^) was an indicator of disease activity. Thus, the role of Treg in IBD requires a more nuanced approach then simple enumeration of T cells bearing classic Treg markers.

It is thought that the development of UC is underpinned by an imbalance between Th17 and Treg cells. UC is associated with a sequestration of immune cells within the gut mucosa, where a pro-inflammatory cytokine environment restricts Treg activity and promotes the continual differentiation and development of a dysregulated Th17 response (37). The generation of Th17 cells requires the expression of RORγt, originally defined as a thymic specific isoform of RORγ (38). In moderate and severe UC patients, the inflammatory response is positively correlated with IL-17 expression in colonic specimens (11).

Effector Th17-like Treg have also been described in the IBD scenario. *Ex vivo* secretion of IL-17A and constitutive expression of the chemokine receptor CCR6 along with RORγt in human effector Treg, suggest that these cells are damaging entities (39). Besides its classical role in regulating Th17 cell migration, CCR6 expression on Treg is an important mediator of their recruitment into inflammatory tissues (40). Thus, CD49f high effector Treg expressing increased level of CCR6 at both the transcriptional and protein levels, are likely to present higher adhesion and migration within the extracellular matrix of the intestinal lamina propria and exacerbates IBD. Additionally, previous studies demonstrate a down regulation of the CD49f receptor on the surface of both CD4^+^ and CD8^+^ conventional T cells after migrating into the intestinal lamina propria of patients with IBD (19), suggesting that CD49f is likely to mediate T cell interactions with extracellular components in the lamina propria of the inflamed human colonic mucosa.

Due to the high Treg plasticity, the ontogenesis of Th17-like Treg is still under debate and some authors suggested that they might represent a transient stage of progenitor cells that can convert into either FoxP3^+^ Treg or Th17 cells under certain inflammatory conditions including UC (13, 14, 41). Interestingly, CD49f may be the only marker commonly found in more than thirty different populations of stem cells, including some hematopoietic stem cells (42). Thus, it is possible that CD49f expression is associated with the preservation of Th17-like Treg cells in a progenitor stage, thus contributing to a pro-inflammatory milieu that leads to the development of IBD. The mechanisms that drive CD49f expression on Treg still remain to be elucidated and it is possible that expression is underpinned by TGF-β-related cytokines. Whilst TGF-β signaling is important for development and maintenance of Treg, it also up-regulates the expression of several types of integrin receptors (42). Our previous findings demonstrate that integrin alpha 6 (CD49f) expression on thymic epithelial progenitor cells is modulated by the members of the TGF-β superfamily of proteins (43).

In conclusion, our findings propose a role for CD49f in modulating Treg cell function and differentiation in humans. Whilst absence of CD49f renders Treg more suppressive and likely to play a vital role in immune homeostasis under normal physiological conditions, high expression of CD49f seems to contribute at least in part to the development of pro-inflammatory effector Treg that correlate with disease activity in UC. Our results highlight the importance of CD49f in modulating Treg mechanisms that guide homeostasis in health and dysfunction in disease.

## ACKNOWLEDGMENTS

We thank the volunteers for donating blood samples for this study. HW was supported by an International Postgraduate Research Scholarship, The University of Queensland, Brisbane, Australia and PhD Top-up Scholarship, QIMR Berghofer Medical Research Institute, Brisbane, Australia. J.J.M is supported by a NHMRC CDF Level 2 Fellowship (1131732). The authors thank the expertise of staff from the Translational Research Institute FlowCore facility and the QIMR Berghofer Analytical facility.

## AUTHOR CONTRIBUTIONS

A.L, J.J.M and M.M.H initiated the project. A.L, H.W, M.M.H and J.J.M designed the methodology. J.B, and S.A provided clinical samples. H.W, K.L, L.C, S.A, A.L conducted experiments. H.W, K.L, J.S, and A.L. analyzed data. M.M.H, C.S, J.J.M, S.L, J.B, A.L. supervised the project. AL, HW, MMH drafted the manuscript. J.J.M is supported by a NHMRC CDF Level 2 Fellowship (1131732) with funding acquisition from The QIMR Berghofer Medical Research Institute, Australia and The Australian Institute of Tropical Health and Medicine (AITHM), Australia. All authors approved the manuscript.

## METHODS

### UC patients and healthy donor specimens

Cryopreserved PBMC from 23 UC patients part of the Mater Hospital IBD biobank, Brisbane, Australia were assessed by flow cytometric analysis (Table 1). Age and gender matched healthy PBMC were included as controls. Blood samples were collected as part of the Mater IBD biobank (Mater HREC approval AM/MML/24730). Patients receiving anti-TNF therapy where excluded from these analysis due to direct effect of TNF-α on Treg (44). Before staining, cryopreserved samples were thawed and incubated in RPMI 1640 containing 10 µg/ml DNAse I (Roche, CH) at 37°C for 1 h to prevent cell clumping and debris. Healthy controls PBMC were freshly isolated from volunteers at QIMR Berghofer for transcriptome, proteomic and functional studies. Ethics approval was obtained from the human research ethics committee QIMR Berghofer, Brisbane, Queensland, Australia (HREC #P2058). In all cases, PBMC were isolated using a Ficoll-Paque Plus (Merck, USA) density gradient centrifugation from blood and a written informed consent was obtained from volunteers.

### Isolation of human T cells

CD4^+^ T cells were separated from healthy controls PBMC using pan T cell isolation kit and magnetic-activated cell sorting (Miltenyi Biotec, USA). Enriched T cells were subsequently stained with CD25-PE mAb and CD25^+^ T cells positively selected using anti-PE magnetic beads and MACS (Miltenyi Biotec, USA). To obtain high purity Treg, CD25^high^ T cells were stained with CD3-APCe780, CD4-BV711 and CD127-BV786 mAbs (BD Biosciences), and zombie aqua live/dead marker (Biolegend, USA) and sorted in FACSAria III (BD Biosciences, USA). The purity of the sorted Treg subsets was assessed by staining the cells with FoxP3-APC mAb using the FoxP3 Fix/Perm Buffer Set for nuclear staining (Biolegend, USA). Over 95% of purity was detected in all cases analyzed. The CD25^-^ fraction was assessed as conv CD4^+^ populations.

### Proteomic sample preparation and LC-MS/MS analysis

Approximately 10^6^ cells were lysed in a SDS-containing buffer for proteomic analysis. Trypsin digestion using the protein co-precipitation method with trypsin and peptide desalting was performed as described (45). Based on micro-BCA assay protein quantification, 0.9 μg of tryptic peptide samples label free shotgun proteomic data were obtained on an Orbitrap Fusion^TM^ Tribrid^TM^ mass spectrometer (Thermo Fisher Scientific, USA) inline coupled to nanoACQUITY ultra performance liquid chromatography system (Waters, USA), using a Symmetry C18, 2G, VM (100Å, 5 μm particle size, 180 μm x 20 mm) trap column and a BEH C18 (130Å, 1.7 μm particle size, 75 μm x 200 mm) analytical column at a flow rate was 3 μl /minute over 175 minutes. The mobile phase consisted of solvent A (0.1% formic acid) and solvent B (100% acetonitrile/0.1% formic acid). Three consecutive linear gradients were used for peptide elution: 5%-9% of solvent B between 3-10 minutes, 9%-26% of solvent B from 10-120 minutes and 26%-40% of solvent B from 120 to 145 minutes. Column cleaning and equilibration was achieved with gradient from 40-80% of solvent B at 145-152 minutes, holding at 80% until 157 minutes and then at 1% until 160 minutes. EASY-Max NG™ ion source (Thermo Fisher Scientific, USA) was used at 1900V and 285°C to ionize the eluted peptides. Xcalibur software (version 3.0.63, Thermo Fisher Scientific, USA) was used with “top speed” mode allowing automatic selection of positively charged peptides (+2 to +7) in a 2 second cycle time.

### Identification of signature proteome of Treg cells

Acquired proteomic data were searched using MaxQuant (Release 1.5.8.3) software (46) against UniProt human reviewed proteome database containing 20,242 entries (downloaded on 25th October 2017). Carbamidomethylation was assessed as the fixed modification while oxidation and N-terminal acetylation were considered as the variable modifications. maxLFQ included in MaxQuant software was used to obtain the normalized label free peptide and protein intensity data. Proteins quantified by at least 2 unique or razor peptides at m-score of > 5 and on > 50% of the samples were selected for further analysis. Missing protein intensity values of the selected proteins were imputed using maximum likelihood estimate (R package) and differential expression analysis was performed using multiple t-test with FDR determination by two-stage linear step-up procedure of Benjamini, Krieger and Yekutieli. In the DE analysis, protein expression data of Treg cells were compared to conv CD4^+^ cells to obtain the log2FC and the statistical significance. DE proteins were defined as log2FC ≥ 1 or ≤ -1 at FDR value of < 0.05. These proteins were further analyzed using IPA core analysis to identify proteins related to cell surface. Each of the proteins annotated as plasma membrane in subcellular localization were individually searched to identify potentially uncharacterized surface proteins in circulating human Treg cells.

### Antibodies and flow cytometry

Fluorescence dye-labeled Abs specific for CD3 (SK7), CD4 (SK3), CD25 (BC96), CD127 (HIL-7R-M21), CD45RA (HI100), CD49f (GoH3), IL-17A (BL168), IFNγ (4S.B3), CTLA4 (L3D10), CD39 (A1), CCR6 (G034E3), and FoxP3 (206D) were purchased from BD Biosciences, eBioscience, and Biolegend, (All, USA). Intranuclear staining for FoxP3 was achieved using a nuclear staining buffer set (Biolegend, USA). Intracellular IL-17A staining was performed using an intracellular staining buffer (BD Biosciences, USA). Viability of cells was defined by LIVE/DEAD™ Fixable Aqua Dead Cell Stain Kit (Thermo Fisher Scientific, USA). Samples were acquired by a BD Fortessa multiparametric flow cytometer (BD Biosciences, USA).

### *In vitro* T cell suppression assay

Suppression of conv CD4^+^ cell proliferation by Treg cells was assessed based on an assay previously optimized for small number of cells (47). Treg cells were sorted as CD49f high or CD49f neg populations. The assay was carried out in a round bottom 96 well plate where 25,000 of conv CD4^+^ cells previously stained with cell trace violet (CTV, Thermo Fisher Scientific, USA) were co-cultured with Treg cells at Treg: conv CD4^+^ cell ratio ranging from 1:1 to 1:16. Four technical replicates were used in each condition and analyzed. To stimulate proliferation, conv CD4^+^ cells were activated with 1μg/ml of soluble anti-human CD3 /OKT3 mAb (Sigma, USA) in the presence of irradiated allogenic PBMC (∼50,000) for five days. Conv CD4^+^ cells stimulated without Treg cells were also included to monitor basal T cell proliferation. Unstimulated conv CD4^+^ cells were used to establish the gating for CTV^-/low^ cells. CTV-based proliferation of conv CD4^+^ cell percentage was assessed with FlowJo software (TreeStar Inc., USA). The percentage of Treg-mediated suppression was calculated as 100−((proliferated cells with Treg/proliferated cells with no Treg) x 100).

### Assessment of cytokine production by Treg

IL-17A and IFNγ cytokines production by enriched CD49f neg and CD49f high Treg was assessed *ex vivo* using intracellular staining. Briefly, 125,000 T regs were plated per well on a 96 well round bottom plate in the presence of 500 IU/ml of human recombinant IL-2 (Novartis, CH) and CD3/28 Dynabeads, using three Dynabeads per one Treg cell (Thermo Fisher Scientific, USA) for activation. Cells were incubated overnight at 37°C. Brefeldin A was added at 10 μg/mL (BD Biosciences, USA) was added for the last 4 hours of incubation. After this period, cells were fixed/ permeabilized with Cytofix/ Cytoperm kit (BD Biosciences, USA) and stained with IL-17A and IFNγ mAbs (Biolegend, USA) for flow cytometric analysis.

### RNA Library Preparation

RNA was purified from approximately 25,000 of FACS-isolated CD49f high and CD49f neg Treg using the Arcturus PicoPure Isolation Kit (Thermo Fisher Scientific, USA). RNA integrity was confirmed on the Agilent 2100 Bioanalyser using the Total RNA Pico Kit (Agilent Technologies, USA). Oligo d(T) captured mRNA was processed for Next Generation Sequencing (NGS) using the NEB Next Ultra II RNA Library Prep Kit for Illumina (New England Biolabs, UK). Quality was assessed on the Agilent 2100 Bioanalyser using the High Sensitivity DNA Kit (Agilent, USA), and quantification was based on the Qubit DNA HS Assay Kit (Thermo Fisher Scientific, USA). Final libraries were pooled and sequenced using a High output single-end 75 cycle (version 2) sequencing kit using the Illumina Nextseq 550 platform (Illumina, USA).

### Bioinformatics analysis of RNA-sequencing data

Reads were trimmed for adapter sequences using Cutadapt (version 1.11) (48) and aligned by STAR (49) (version 2.5.2a) to the GRCh37 assembly using the gene, transcript, and exon features of Ensembl (release 70) gene model. Expression was estimated through RSEM (version 1.2.30) (50). Transcripts with zero read counts across all samples were removed prior to analysis. Normalization of read counts was performed by dividing million reads mapped to generate counts per million (CPM), followed by the trimmed mean of M-values (TMM) method from the edgeR package (version 3.32.0). For the differential expression analyses, the glmFit function was adapted to fit a negative binomial generalized log-linear model to the read counts for each transcript. Using the glmLRT function, we conducted transcript wise likelihood ratio tests for each comparison. Log2 transformed, normalized read counts were used for heatmaps and PCA. Hierarchical clustering of genes and samples in the heatmaps was performed with average linkage clustering on the 1- Spearman correlation coefficient dissimilarity matrix of all DE transcripts with a FDR <0.05 or as stated in text.

### Gene set enrichment and pathway analysis

Gene set enrichment analysis was performed using GSEA from Broad Institute (51). P-values were generated form 1000 gene set permutations, excluding gene sets with more than 3000 genes or less than 5 genes against custom made gene sets (GSE107011) (52, 53) and Broads Hallmark database. In addition, IPA was used with the default settings to identity canonical pathways of DE genes (FDR < 0.05) transcripts.

### Quantification and statistical analysis

All values are expressed as mean ± SEM, unless otherwise specified. Statistical analyses were performed using GraphPad Prism v7.02 (GraphPad Software, USA), with the appropriate tests utilized. A p value < 0.05 was considered statistically significant.

### Data and Code Availability

RNA-Seq data reported in this paper is available on the GEO repository with accession number GSE161077.

The mass spectrometry proteomics data have been deposited on the ProteomeXchange Consortium via the PRIDE (54) partner repository with the dataset identifier PXD022095.

## SUPPLEMENTARY FIGURE LEGENDS

**Supplementary figure 1. Comparative proteomic analysis of circulating regulatory T cells and conventional CD4^+^ T cells.**

(a) Percentage of missing maxLFQ protein intensity values in three donors analyzed.

(b) Data mining steps used in selection of constantly identified high confident proteins for DE analysis between conv CD4^+^ and Treg cells in three donors analyzed.

(c) Number of proteins quantified at different unique and razor peptide numbers.

(d) Protein intensity distribution patterns in three donors analyzed. Box plots comprise of central lines and boxes representing mean values and 95% confidence intervals, respectively. Whiskers representing 2.5 to 97.5 percentiles.

(e) Heatmap analysis of differentially expressed proteins from comparing Treg and conv CD4^+^ cell. Each column represents individual donors across proteins differentially expressed between Treg versus conv CD4^+^ cell (n=3; FDR < 0.01).

(f) Box plots displaying the covariance (CV%) of intensities between conv CD4^+^ and Treg cells. Boxplots comprise of central lines and boxes representing mean values and 95% confidence intervals, respectively. Whiskers are 2.5 to 97.5 percentiles.

**Supplementary figure 2. Regulatory T cell-mediated suppression and pro-inflammatory cytokines production.**

(a) Graph showing the fraction of proliferative conv CD4^+^ cells cultured in the presence or absence of total Treg at various Treg: conv CD4^+^ cell ratios. Data represents one donor analyzed and includes 3 technical replicates. *p<0.05, **p<0.01 and ***p<0.001. Non-parametric paired t-test. Bars represent standard error of mean.

(b) Representative FACS dot-plot showing IL-17A expression on total Treg following overnight activation in the presence of CD3/CD28 Dynabeads and recombinant IL-2 during five days of *in vitro* culture.

(c) Fraction of IFNγ^+^ cells in CD49f neg and CD49f high Treg. Data was obtained from four healthy controls analyzed.

(d) Representative FACS dot-plot showing IL-17A and IFNγ co-expression on CD49f high Treg.

**Supplementary figure 3. Genes modulated in CD49f negative versus CD49f high regulatory T cells.**

(a) Pie chart showing total genes analyzed and DE genes between CD49 high and CD49 neg Treg using RNA-Seq data across multiple donors.

(b) Number of genes over-expressed and under-expressed from total DE genes between CD49f high and CD49 neg Treg using RNA-Seq data across multiple donors.

**Supplementary figure 4. Modulation of immune markers in CD4^+^ T cell subsets.**

Flow cytometry generated dot-plots quantifying MFI expression of CD4, CD25, CD39, CTLA4, CCR6 and CD25 in CD4^+^ T cell subsets in healthy PBMC. Cells were classified as effector Treg (CD25^+^CD127^-^FoxP3^high^CD45RA^-)^, resting Treg (CD25^+^CD127^-^ FoxP3^low^CD45RA^+^) and FoxP3^+^ non Treg cells (CD25^+^CD127^-^FoxP3^low^CD45RA^-^). Each symbol represents an individual donor analyzed. *n.s.* = non-significant, P*<0.05, P**<0.01, P***<0.0005, P****<0.0001. Non-parametric one-way ANOVA test with Bonferroni correction.

**Supplementary figure 5. Comparison of conventional CD4^+^ T cells and Treg in healthy controls and in UC patients.**

Frequencies of Treg in PBMC from patients with active UC (n = 10), non-active UC (n = 13) and healthy controls (HC, n = 10).

(a) Proportion of CD4^+^ T cells in total T cells. *n.s.* = non-significant, P*<0.05. Non-parametric one-way ANOVA test with Bonferroni correction. Bars represent standard error of mean.

(b) Proportion of Treg cells in total CD4^+^ T cells. *n.s.* = non-significant. Non-parametric one-way ANOVA test with Bonferroni correction. Bars represent standard error of mean.

(c) Proportion of CD49f high cells in resting Treg. *n.s.* = non-significant. P*<0.05. Non-parametric one-way ANOVA test with Bonferroni correction. Bars represent standard error of mean.

(d) Proportion of CD49f neg cells in resting Treg. *n.s.* = non-significant. Non-parametric one-way ANOVA test with Bonferroni correction. Bars represent standard error of mean.

(e) Resting Treg CD49f index, representing the ratio of CD49f high over CD49f neg resting Treg. *n.s.* = non-significant. Non-parametric one-way ANOVA test with Bonferroni correction. Bars represent standard error of mean.

## SUPPLEMENTARY TABLES

Supplementary table 1. Surface membrane proteins identified from proteomic study.

Supplementary table 2. Significant DE proteins between regulatory T cells and conventional CD4^+^ T cells.

Supplementary table 3. Significant DE genes between CD49f high and negative Treg identified from transcriptomic analysis.

