## Supplementary Figures for "Expression of CD49f defines subsets of human regulatory T cells with divergent transcriptional landscape and function that correlate with ulcerative colitis disease activity"

### Supplementary figure 1

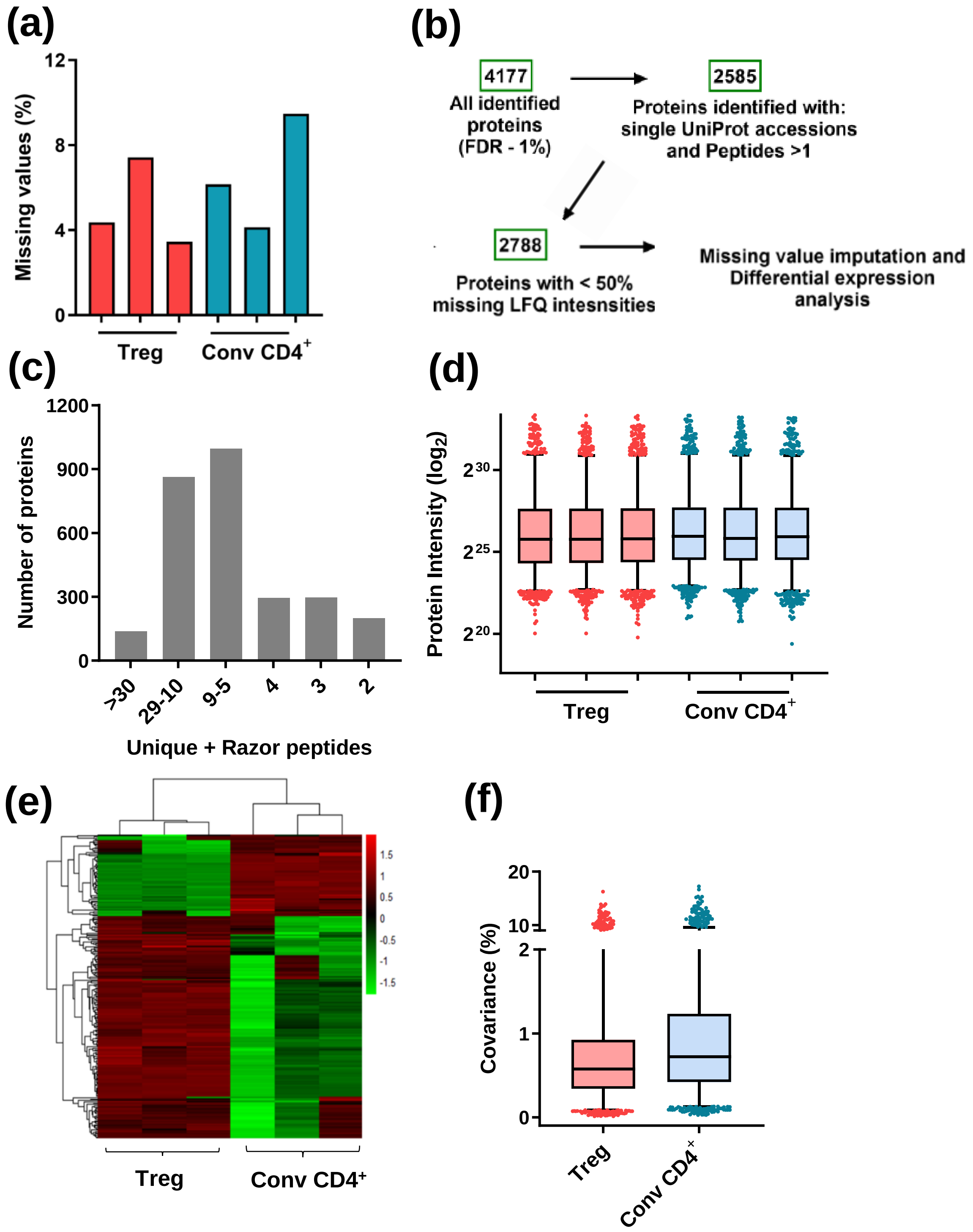

### Supplementary figure 2

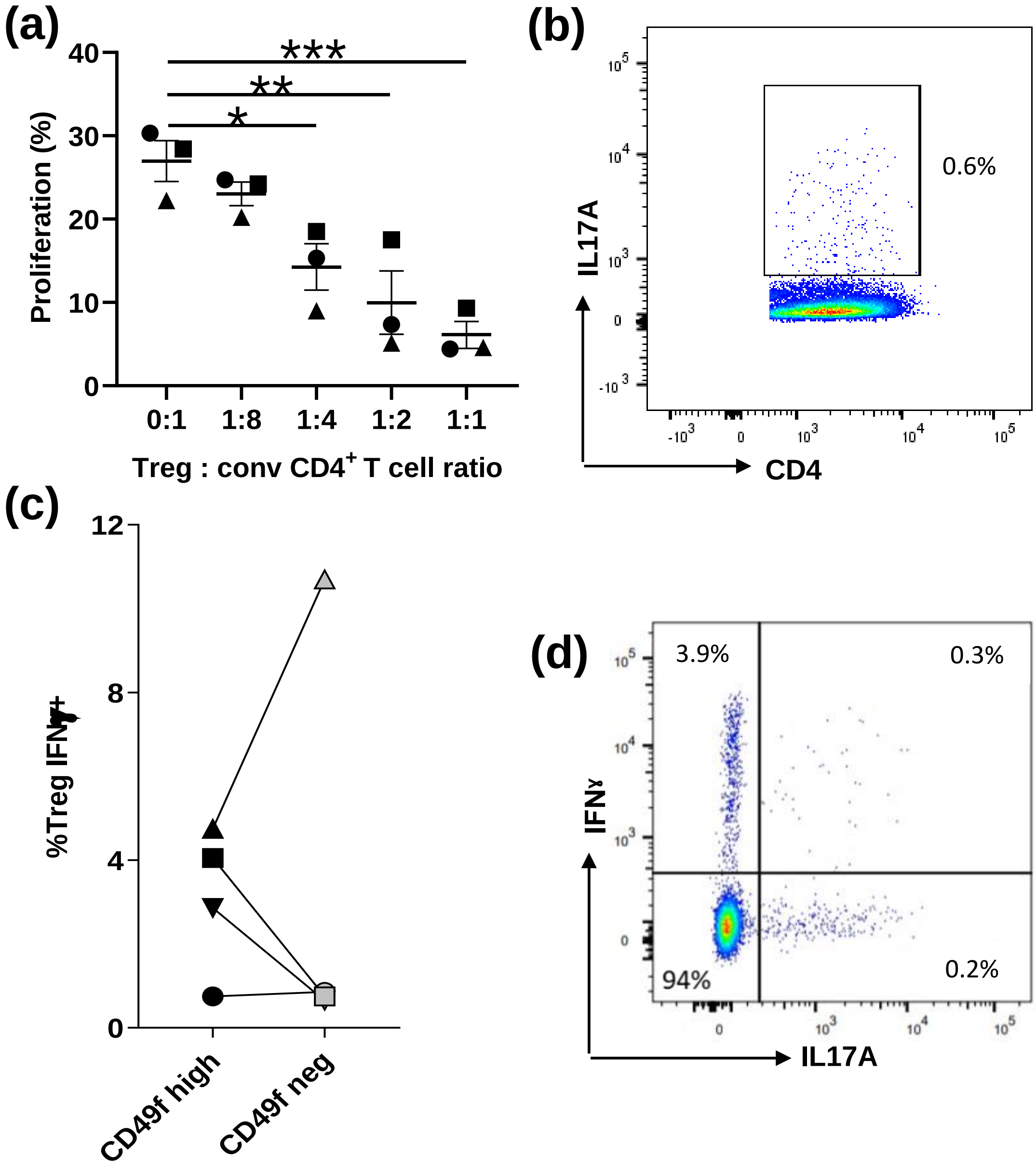

### Supplementary figure 3

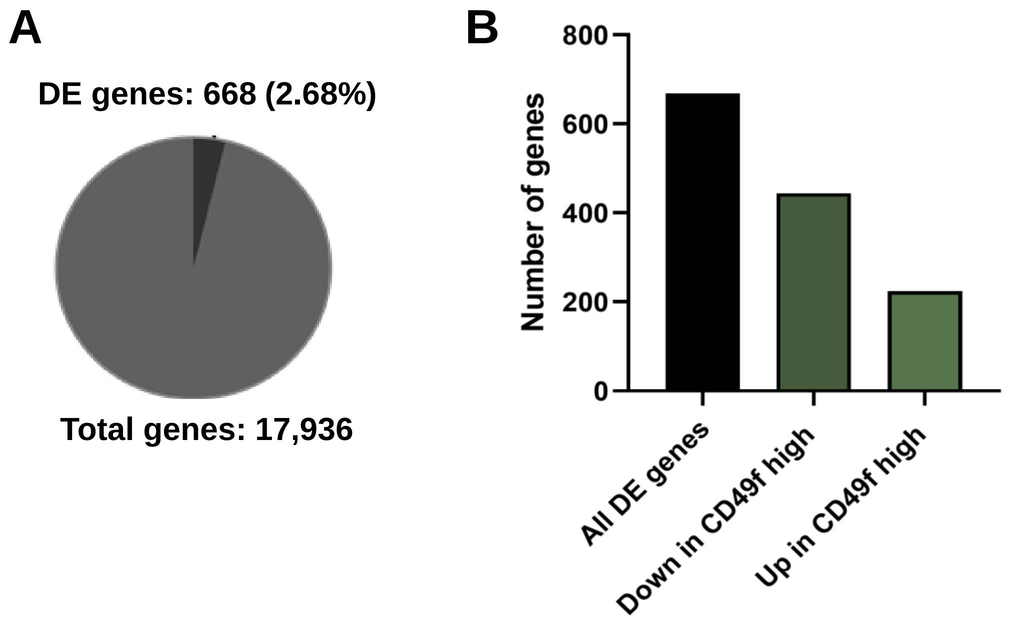

### Supplementary figure 4

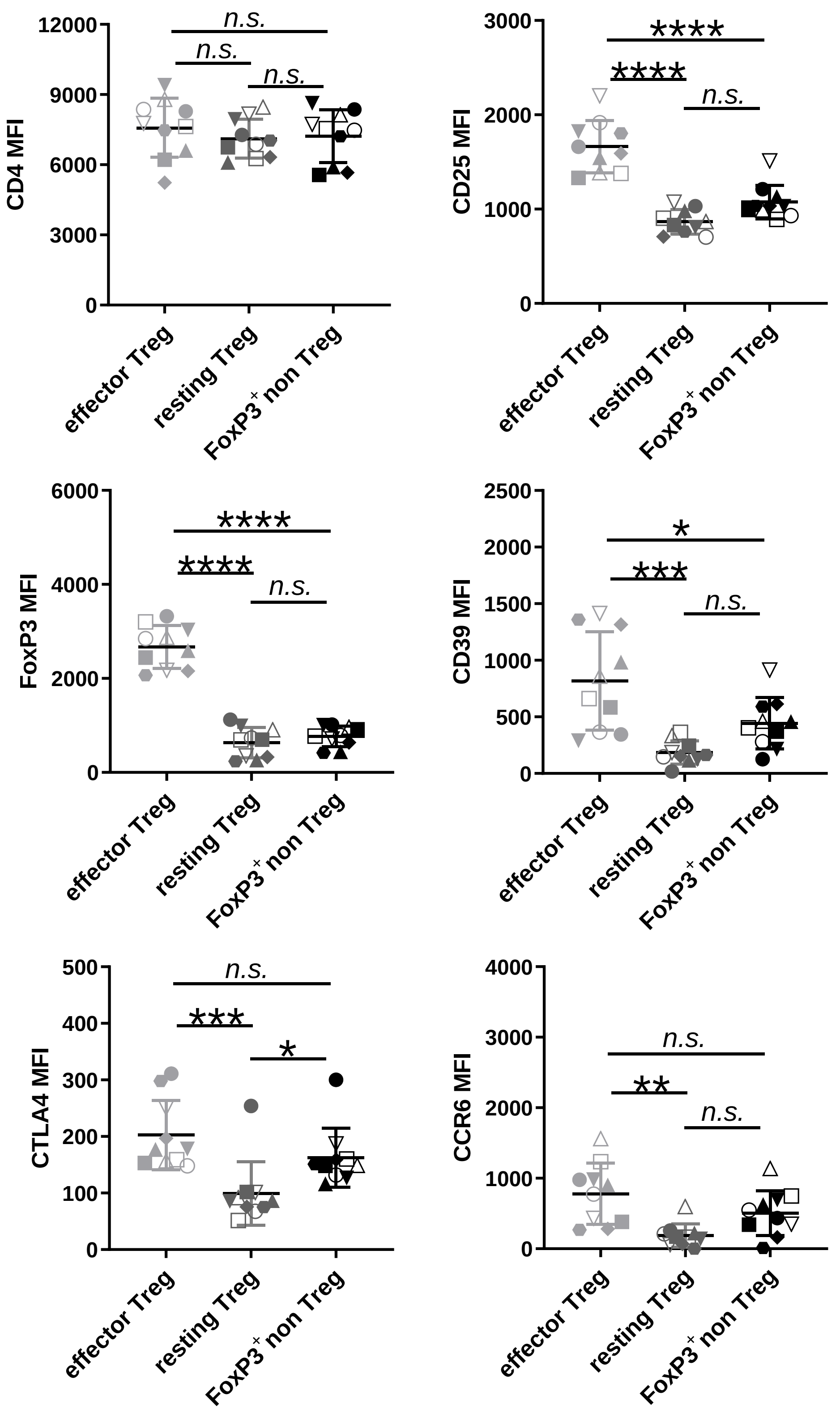

### Supplementary figure 5

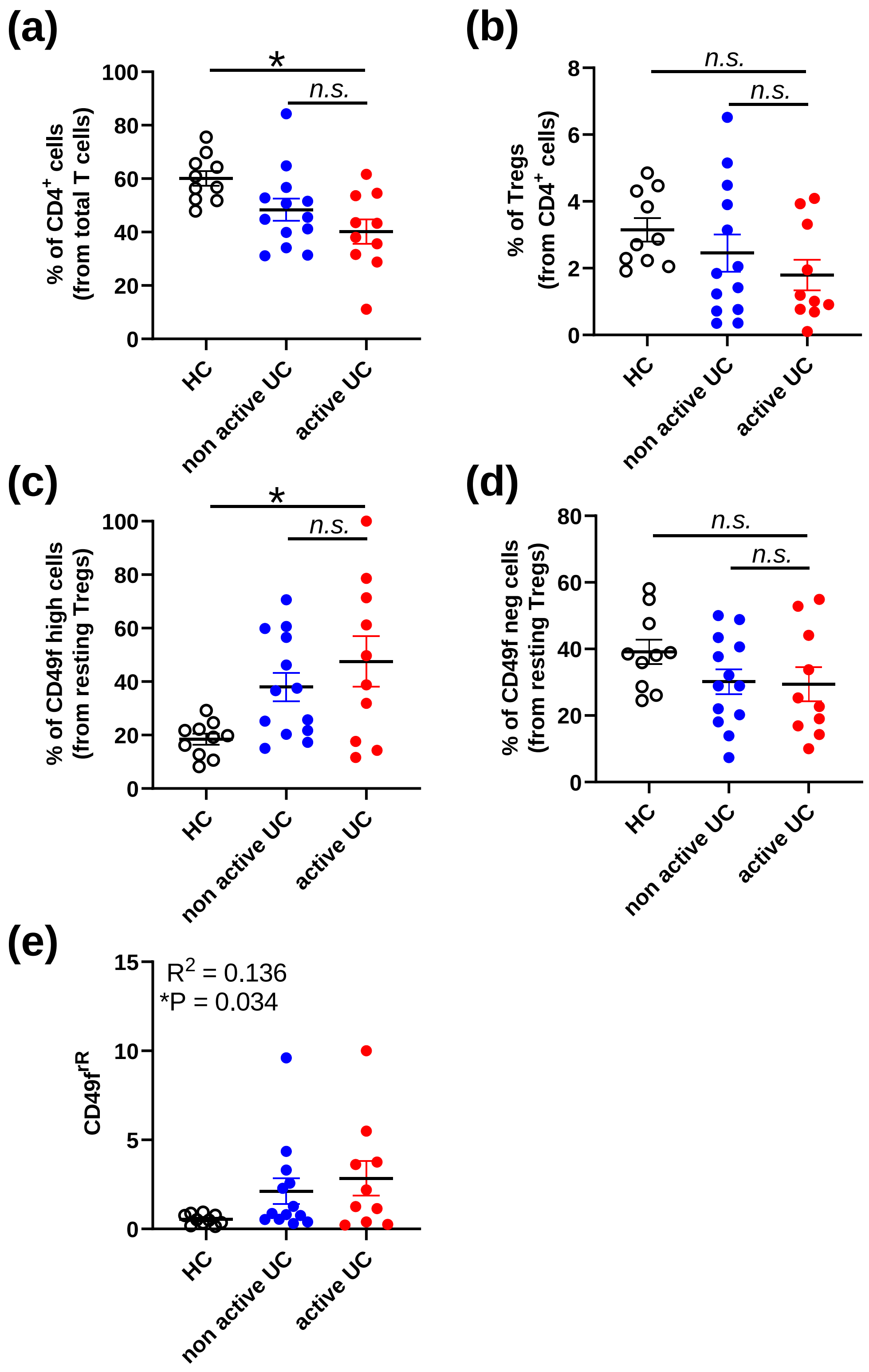
